# Ratiometric NAD^+^ sensors reveal subcellular NAD^+^ modulators

**DOI:** 10.1101/2022.05.20.491061

**Authors:** Liuqing Chen, Meiting Chen, Mupeng Luo, Yong Li, Bagen Liao, Min Hu, Qiuliyang Yu

**Affiliations:** Shenzhen Institute of Advanced Technology, Chinese Academy of Sciences, 518055 Shenzhen, China; Shenzhen Key Laboratory for the Intelligent Microbial Manufacturing of Medicines, 518055 Shenzhen, China; Department of Sports Medicine, Guangzhou Sport University, 510150 Guangzhou, China

## Abstract

Mapping NAD^+^ dynamics in live cells and human are essential for translating NAD^+^ interventions into effective therapies. Yet genetically encoded NAD^+^ sensors with better specificity and pH-resistance are still needed for cost-effective monitoring of subcellular and clinical NAD^+^. We introduce multicolor, resonance energy transfer-based NAD^+^ sensors that cover nano- to milli-molar concentration ranges for clinical NAD^+^ measurement and subcellular NAD^+^ visualization. The sensors captured the blood NAD^+^ increase induced by NMN supplementation and revealed the distinct subcellular effects of NAD^+^ supplements and modulators. Then, the sensors enabled high-throughput screenings for mitochondrial and nucleic NAD^+^ modulators and identified α-GPC, a cognition-related metabolite, induces NAD^+^ redistribution from mitochondria to nucleus relative to the total adenine nucleotides, which was confirmed by NAD^+^ FRET microscopy.

## INTRODUCTION

Nicotinamide adenine dinucleotide (NAD^+^) functions as a coenzyme in over 300 oxidation-reduction reactions, making it central for energy metabolism. NAD^+^ is also a cofactor and substrate of multiple non-redox enzymes^1^ that are crucial in maintaining cell functions including sirtuins, PARPs, CD38 and SARM1. Recently, low NAD^+^ levels have been linked to multiple disease states, therefore NAD^+^-boosting approaches have been proposed as a potential therapeutic strategy against various diseases^2–4^. In fact, increasing evidences suggest that NAD^+^-boosting strategies can improve physiological and metabolic health in rodent models of ageing, obesity, and diabetes^5,6^, and recent clinical studies suggested a clear association between NAD^+^ and physiological status during human aging^7^ and disease progression^8^.

Measuring NAD^+^ levels in clinical samples, live cells, and animals are essential for understanding the NAD^+^ metabolic network and ultimately translating the NAD^+^-boosting therapies to human. NAD^+^ sensors based on semisynthetic and genetically encoded protein constructs have been introduced to monitor NAD^+^ levels under various settings^9–11^. Although the semisynthetic NAD^+^ sensor is suitable for point-of-care applications^10^, the limited cell permeability of the sensor’s synthetic fluorophore hinders its application in live cells and animals^12^. Genetically encoded NAD^+^ sensors based on circular-permuted fluorescent proteins can work in live cells with good spatiotemporal resolutions, however the pH-sensitivity and substrate specificity to discriminate against NADH, NR, NMN, and AXP can still be improved for the reported NAD^+^ sensors^9,12,13^.

Here, we developed a set of genetically encoded multicolor NAD^+^ sensors (NS) named as NS-Goji, NS-Olive, and NS-Grapefruit using Bioluminescent and Fluorescent Energy Transfer (BRET and FRET). We further used the sensors to visualize subcellular NAD^+^ dynamics in live cells and to screen for organelle-specific NAD^+^- modulating compounds. In the screening, we have identified L-Alpha glycerylphosphorylcholine (α-GPC) as a modulator of NAD^+^ distribution in mitochondria and nucleus relative to the total adenine nucleotide.

## RESULTS

### Genetically encoded BRET and FRET sensors for NAD^+^

We designed the genetically encoded BRET NAD^+^ sensors using a NAD^+^-sensitive domain, a circularly permuted NanoLuc luciferase (cpNLuc) and a fluorescent protein (Fig. 1A). We used the Pro5-to-Pro317 domain of DNA ligase from *Enterococcus faecalis* V583 as the NAD^+^-sensitive domain (denoted as *Ef*LigA) for this domain is known to undergo considerable conformational changes^14^ upon binding to NAD^+^ and has no reported affinity towards NADH, making it advantageous for developing a NAD^+^-specific sensor with minimal response to NADH. Then, fluorescent protein mScarlet and luciferase cpNLuc were fused to N and C termini of the NAD^+^-sensing domain to form the BRET NAD^+^ sensor scaffold. We then performed rounds of protein engineering to optimize its response towards NAD^+^ (Fig. 1B). Based on the 3D structures of *Ef*LigA (PDB ID: 1ta8), disrupting domain interactions in its apo state may lead to more separated N and C termini which should enlarge the sensor’s signal response. Hence, we targeted residues R68 and R163 that potentially participate in the domain interaction within the apo-*Ef*LigA (Fig. S1) and obtained a sensor mutant (R68A/R163D) with a slightly improved signal response toward NAD^+^ (represented by R_max_/R_min_ = 1.09, Fig. 1C). We then optimized the linker region between *Ef*LigA and mScarlet (L1, STGPLT) and that between *Ef*LigA and cpNLuc (L2, GLSGD) by screening amino acid residues and by reducing the linker length. After screening 175 mutants, the R_max_/R_min_ was improved to 1.89 with linker L1 changed from STGPLT to STLT, and L2 from GLSGD to VD (Fig. 1B, C). To further enlarge the R_max_/R_min_, shorten the fluorophore’s maturation time, and improve the sensitivity, we performed saturated mutagenesis on R68 and R163, replaced mScarlet with mScarlet-I and mutated residues in the NAD^+^-binding pocket of *Ef*LigA (Fig. 1B). Through these processes, four versions of the BRET NAD^+^ sensor (denoted as NS-Goji 1.0, 1.1, 1.2 and 1.3) with satisfactory dynamic ranges have been developed to cover the different concentration windows of NAD^+^ (Fig. 1D). NS-Goji 1.0 carries the mutation R68N/R163D on *Ef*LigA domain and features a c50 of 235.7 ± 7.9 μM. An additional mutation Y42W introduced on *Ef*LigA gave rise to NS-Goji 1.1 with a c50 of 3.1 ± 0.3 mM. Mutation R68N, R163D, D91Q and Y227F on *Ef*LigA yielded NS-Goji 1.2 with a c50 of 7.0 ± 0.1 μM. Mutation R68N, R163D, D91K and Y227F further reduced the c50 to 47.4 ± 0.9 nM (NS-Goji 1.3). The sensors showed a high specificity towards NAD^+^ over other structurally similar molecules and did not show apparent response towards NAD^+^ precursors such as NR, NMN and NRH (Fig. 1E). The sensor is insensitive toward physiological pH changes and works properly in the presence of 0.5 mM NMN or NRH (Fig. S2). The performance of these sensors is summarized in Table S1.

**Fig.1.**
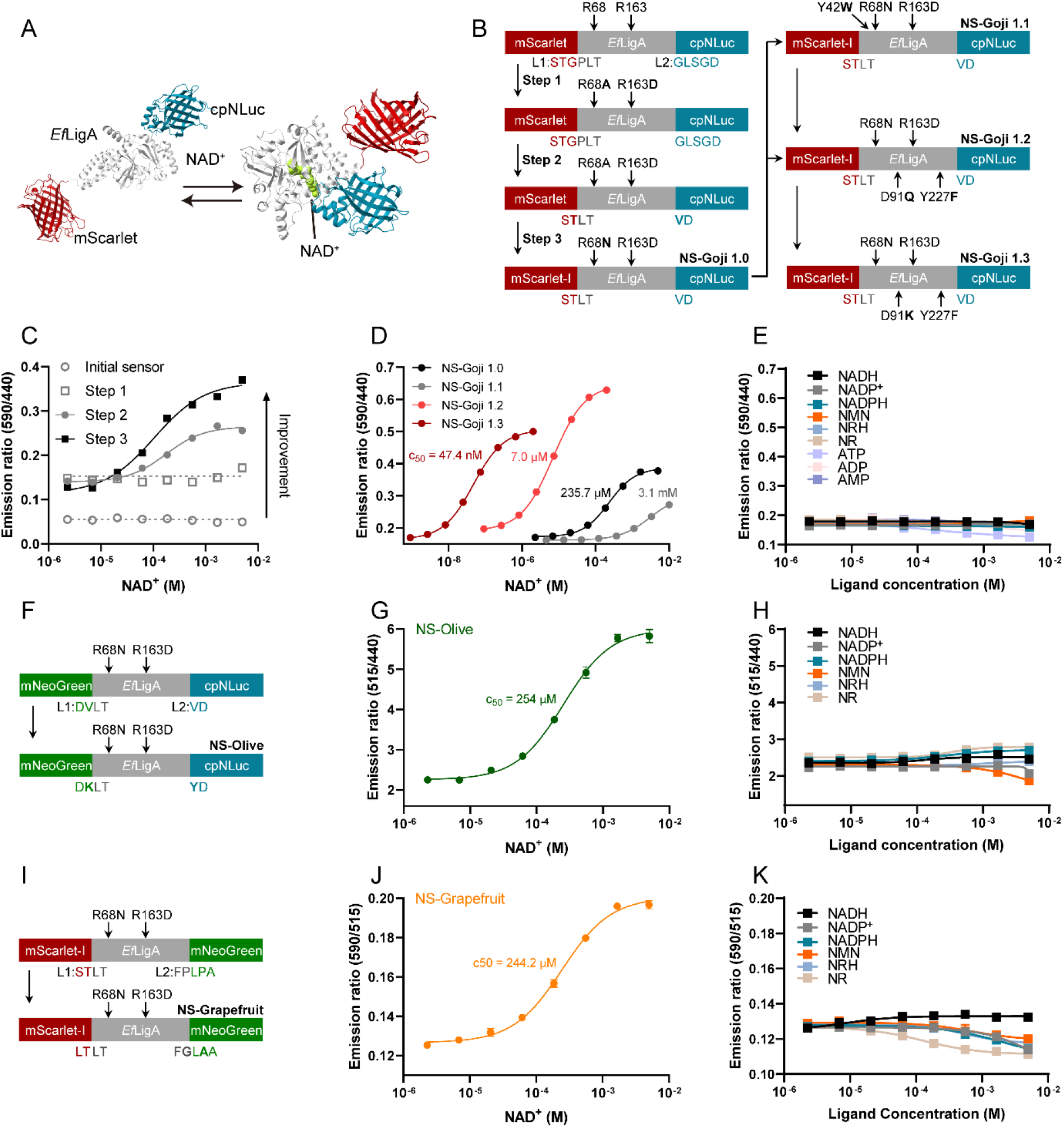
Design and optimization of NAD^+^ sensor. (A) Sensor design: The sensor consists of cpNLuc, *Ef*LigA and mScarlet. The binding of NAD^+^ induces conformational change in *Ef*LigA that pulls cpNLuc and mScarlet in proximity to enable BRET. (B) Workflow of NS-Goji optimization to improve dynamic range and sensitivity. (C) Signal response of NAD^+^ sensor during optimization. (D) Titration of NS-Goji 1.0, NS-Goji 1.1, NS-Goji 1.2, and NS-Goji 1.3 with NAD^+^. (E) Titration of NS-Goji 1.0 with NADH, NADP^+^, NADPH, NMN, NRH, NR, ATP, ADP, and AMP. (F) Workflow of NS-Olive optimization. (G) Titration of NS-Olive with NAD^+^. (H) Titration of NS-Olive with NADH, NADP^+^, NADPH, NMN, NRH and NR. (I) Workflow of NS-Grapefruit optimization. (J) Titration of NS-Grapefruit with NAD^+^. (K) Titration of NS-Grapefruit with NADH, NADP^+^, NADPH, NMN, NRH and NR. Values are given as mean ± SD of three independent measurements.

As organelles have distinct NAD^+^ metabolism, multicolor NAD^+^ sensors can help to simultaneously depict the dynamic regulation of compartmentalized NAD^+^ in live cells. Hence, in addition to the red NS-Goji, we developed a green version of the sensor by replacing the mScarlet-I with mNeoGreen in the sensor scaffold (Fig. 1F). Subsequent screening of site-saturation mutagenesis library on the sensor’s linker region yielded a green BRET NAD^+^ sensor (denoted as NS-Olive) with a c50 of 254 ± 11.5 μM (Fig. 1G) and a signal response of 2.6-fold (Table S1). NS-Olive showed high specificity toward NAD^+^ (Fig. 1H). However, in consistence with the property of mNeoGreen, NS-Olive is slightly affected by pH (Fig. S3) but works properly with the presence of NMN and NRH (Fig. S3).

To facilitate fluorescent imaging of NAD^+^ dynamics in live cells, we further developed a FRET NAD^+^-sensor by replacing the energy donor cpNLuc in NS-Goji 1.0 with mNeoGreen (Fig. 1I). For optimization, we performed site-saturation mutagenesis and linker optimization and identified a FRET sensor with a c50 of 244.2 ± 14.9 μM (Fig. 1J) and a signal response of 1.6-fold (denoted as NS-Grapefruit, Table S1). NS-Grapefruit also showed high specificity toward NAD^+^ (Fig. 1K). Due to the use of mNeoGreen, NS-Grapefruit is slightly affected by pH as well (Fig. S4).

### NS-Goji measures NAD^+^ in human blood samples

As blood NAD^+^ level is a key parameter for assessing the effect of NAD^+^-regulating dietary supplements in several clinical studies^7,15,16^, we tested if our NAD^+^ BRET sensor can capture the increase of NAD^+^ in venous blood induced by supplementing oral NMN. Venous blood samples were obtained from a placebo-controlled, double-blinded clinical study that evaluates the effect of NMN (Registration number: ChiCTR2000040222). Samples from control (Placebo, n = 8) and intervention group (Oral administration of 500 mg NMN for 1 month, n = 8) were lysed and measured using NS-Goji 1.3 (Fig. 2A) for its high sensitivity and pH-resistance (Fig. 2B, C). In parallel, the samples were measured using HPLC-MS as reference. The results obtained by NS-Goji 1.3 and HPLC-MS demonstrated a high level of agreement with a Pearson’s r = 0.980 (Fig. 2D, F), demonstrating that NS-Goji 1.3 can serve for rapid measurement of NAD^+^ levels in clinical samples. The result also confirmed that oral administration of 500 mg NMN for 1 month significantly (*p* < 0.001) increased the NAD^+^ level of venous blood (Fig. 2E).

**Fig.2.**
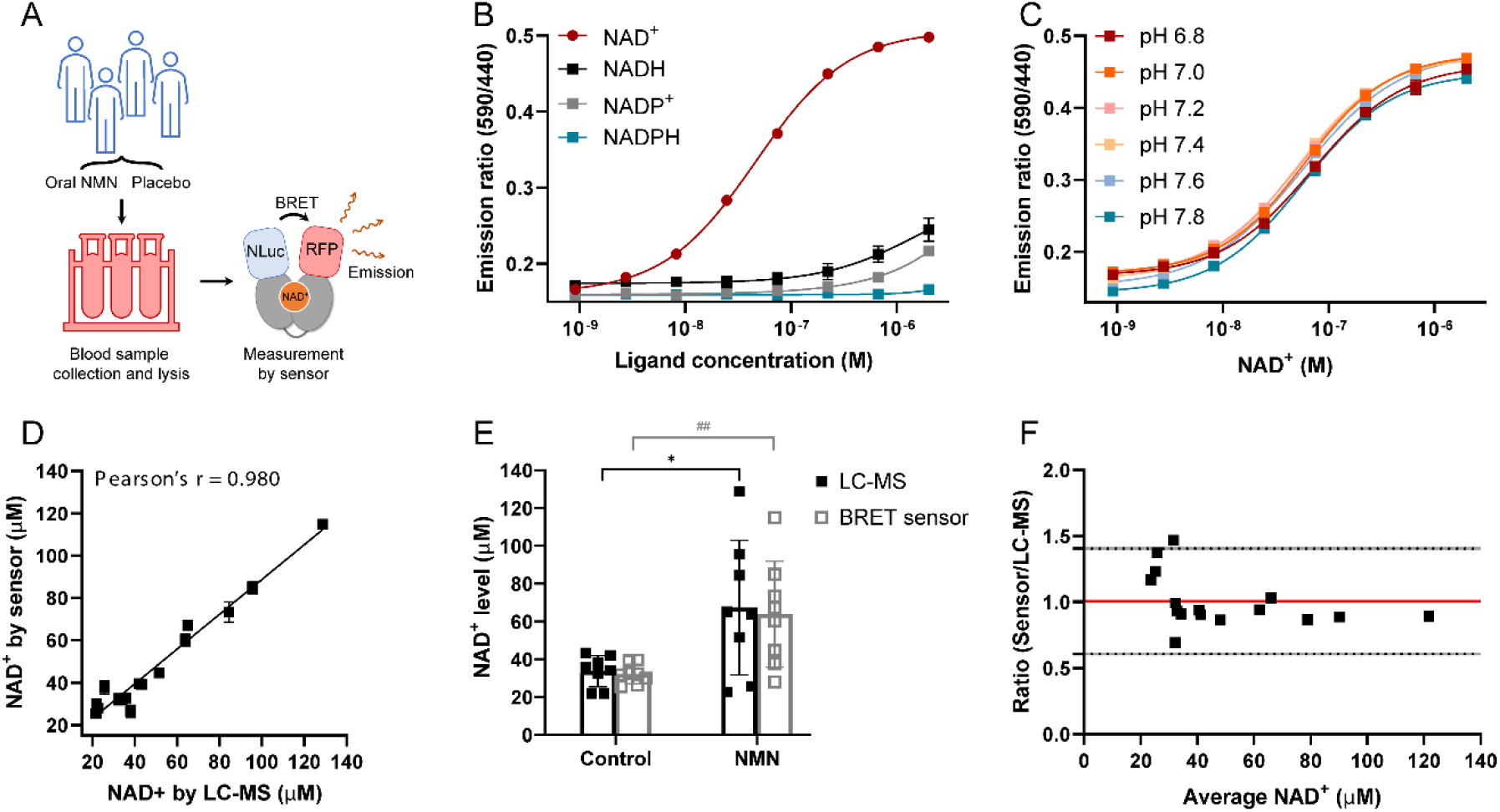
Blood NAD^+^ measurement by NS-Goji 1.3. (A) Workflow of NAD^+^ measurement in clinical samples by NS-Goji 1.3. (B) Titration of NS-Goji 1.3 with NAD^+^, NADH, NADP^+^, and NADPH. (C) Titration of NS-Goji 1.3 with NAD^+^ at various pH. (D) Measurement of NAD^+^ in human blood samples. Sensor results are plotted against LC-MS results. (E) Oral administration of NMN significantly increases NAD^+^ in whole blood (n = 8). (F) Bland-Altman analysis for whole blood NAD^+^ measurement by sensor and LC-MS. In (B), (C), and (D), error bars represent SD of three independent measurements. In (E), values are given as mean ± SD from eight independent samples. Statistical significance was calculated by Student’s t test. *, *P* < 0.05; ^##^, *P* < 0.01

### NS-Olive reveals distinct subcellular effects of NAD^+^-modulating compounds

Unbalanced NAD^+^ metabolism is involved in various age-related diseases. Pharmacological modulation of cellular NAD^+^ holds the promise of intervening a wide range of degenerative diseases^3,17–19^. As NAD^+^ metabolism is compartmentalized and elaborately controlled^20^, understanding the subcellular effects of NAD^+^-modulating strategies are important for evaluating their pharmacological functions. To measure the subcellular NAD^+^, we developed HEK 293T cell lines with stable expressions of NS-Olive in cytoplasm, mitochondria (Mito-NS-Olive), and nucleus (Nuc-NS-Olive) using organelle-targeting peptides (Fig. 3A). The sensor then reports subcellular NAD^+^ levels as ratios of emission intensities at 515 nm and 440 nm measured by a bioluminescent plate reader.

**Fig.3.**
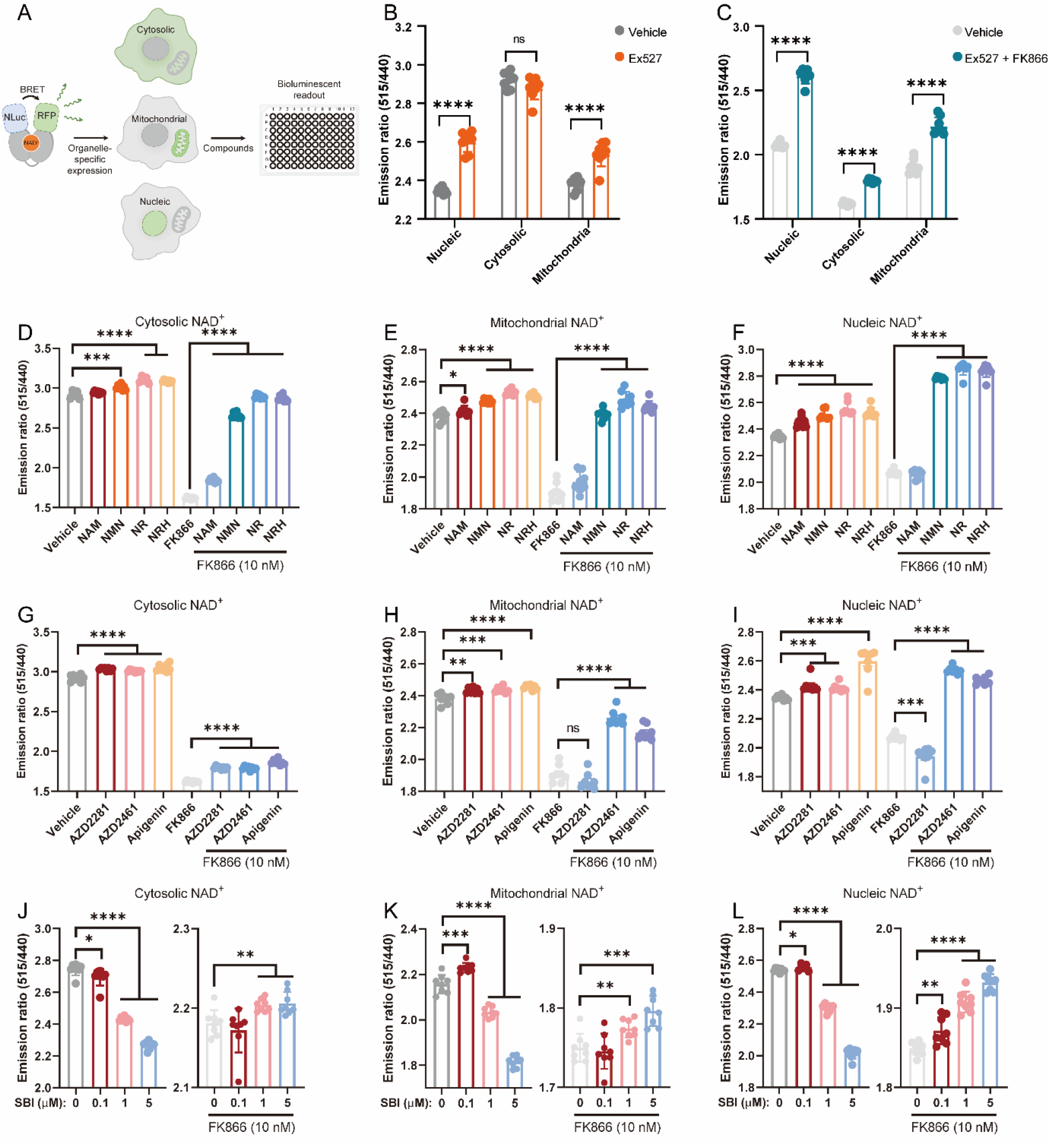
NS-Olive reports compartmentalized NAD^+^ metabolism in live cells. (A) Workflow of measuring NAD^+^ in live cells using organelle specific NS-Olive. (B, and C) Effects of EX527 (100 μM) on cytosolic, mitochondrial, and nucleic NAD^+^ with and without FK866 (10 nM) measured by NS-Olive, n ≥ 6. (D, E and F) Effects of NAD^+^ precursors (500 μM) on cytosolic, mitochondrial, and nucleic NAD^+^ with and without FK866 (10 nM), n ≥ 6. (G, H and I) Effects of apigenin (100 μM), AZD2281 (1 μM), and AZD2461 (1 μM) on cytosolic, mitochondrial, and nucleic NAD^+^ with and without FK866 (10 nM), n ≥ 6. (J, K and L) Effects of NAMPT activator SBI-797812 with and without FK866 (10 nM) on cytosolic, mitochondrial, and nucleic NAD^+^, n ≥ 6. All statistical significance was calculated by Student’s t test. All data are represented as mean ± SD. *, *P* < 0.05; **, *P* < 0.01; ***, *P* < 0.001; ****, *P* < 0.0001.

We first verified the localization of NS-Olive by imaging the sensor’s fluorescent protein in parallel with organelle-specific dyes. The results indicated a good overlap of NS-Olive with mitochondria- and nucleus-specific staining (Fig. S5). We then tested if the compartmentalized NS-Olive can report NAD^+^ changes in subcellular structures. Compound Ex527 preferentially inhibits Sirtuin 1 (IC50 = 38 nM) and has much lower affinity towards Sirtuin 2 (IC50 = 19.6 μM) and Sirtuin 3 (IC50 = 48.7 μM)^21^. Sirtuin 1 locates primarily in nucleus, Sirtuin 2 in cytosol and nucleus, while Sirtuin 3 in mitochondria^22^. We treated the cells with Ex527 and observed that the compound indeed significantly increased the nucleic NAD^+^ (Fig. 3B), but to a less extend the cytosolic and mitochondrial NAD^+^ (Fig. 3B), which is consistent with a previous report^9^. Notably, in the case of FK866-induced NAD^+^ deficiency, Ex527 restored the nucleic NAD^+^ to a level similar to the unchallenged cells (Fig. 3C) but did not show as strong effects on cytosolic nor mitochondrial NAD^+^ (Fig. 3C).

We then tested the effects of various NAD^+^-modulating conditions. We showed that NAD^+^ precursors such as NMN, NR and NRH significantly increased the NAD^+^ levels in cytosol, mitochondria, and nucleus of HEK 293T cells (Fig. 3D-F). In NAD^+^-depleted cells (treated by FK866), NMN, NR and NRH restored the NAD^+^ level across the measured subcellular structures with the strongest effect observed in nucleus (Fig. 3F). Meanwhile, Nam rescued the NAD^+^ depletion in cytosol but not in mitochondria nor nucleus (Fig. 3D-F), indicating a marked difference in the NAD^+^ supplementing abilities of the various precursors.

Other NAD^+^-modulating compounds have been evaluated as well. PARP inhibitors AZD2281 and AZD2461 both increased the NAD^+^ across the measured subcellular compartments of HEK 293T cells (Fig. 3G-I). For NAD^+^-depleted cells, AZD2281 and AZD2461 increased the cytosolic NAD^+^, but AZD2461 demonstrated a much stronger boosting effect on mitochondrial and nucleic NAD^+^ compared to AZD2281, which may be explained by AZD2461’s better resistance to drug transporters and its higher intracellular concentration^23^. Apigenin is a known CD38 inhibitor with wide-range effects on pathways^24^ including TRAIL, PI3K/Akt/Mtor, and MAPK signaling. Our NAD^+^ reporter cells indicated that apigenin significantly increases cytosolic, mitochondrial, and nucleic NAD^+^ (Fig. 3G-I). Apigenin also rescued the FK866-induced NAD^+^ depletion with a greater effect on mitochondria and nucleus than on cytosol. Recently reported molecule SBI-797812 features concentration-dependent dimorphism on NAMPT activation^25^. SBI-797812 reportedly activates NAMPT but has inhibitory effects at high concentrations. We hence tested the response of subcellular NAD^+^ towards SBI-797812 at different concentrations (Fig. 3J-L). In mitochondria and nucleus, SBI-797812 increases NAD^+^ at 0.1 μM but depletes NAD^+^ at 1 and 5 μM (Fig. 3J-L). However, in cytosol, no obvious NAD^+^-boosting effects were observed at the tested concentrations (Fig. 3J). Interestingly, when NAMPT inhibitor FK866 is present, SBI-797812 demonstrates universal NAD^+^-restoring effects with the strongest increase observed in nucleus. Collectively, we showed that the BRET NAD^+^ sensors are viable tools for measuring subcellular NAD^+^. The sensor further revealed that different modulating strategies have distinct effects on the compartmentalized NAD^+^.

### High-throughput screening for novel NAD^+^-modulating compounds

Compartmentalized NAD^+^ regulates various cell functions, yet there is a lack of tool compounds for the organelle-specific modulation of NAD^+^. To screen for compounds that regulate subcellular NAD^+^ levels, we used HEK 293T cells that specifically express NS-Olive in mitochondria and nucleus to screen a total number of 1144 compounds from the APExBIO FDA-approved drug library (Fig. 4A). We used the cellular bioluminescent intensity to indicate cell viability and the emission ratio of 515 nm and 440 nm to indicate subcellular NAD^+^ levels. We first eliminated compounds that decreased cell viability to lower than 70% compared to the control and then selected compounds that increased mitochondrial or nucleic NAD^+^ (Fig. 4B-E). For both mitochondria and nucleus, we listed the top 20 compounds that increased NAD^+^ in respective organelles (Table S2 and S3), among which 11 compounds overlap (Fig. 4F).

**Fig.4.**
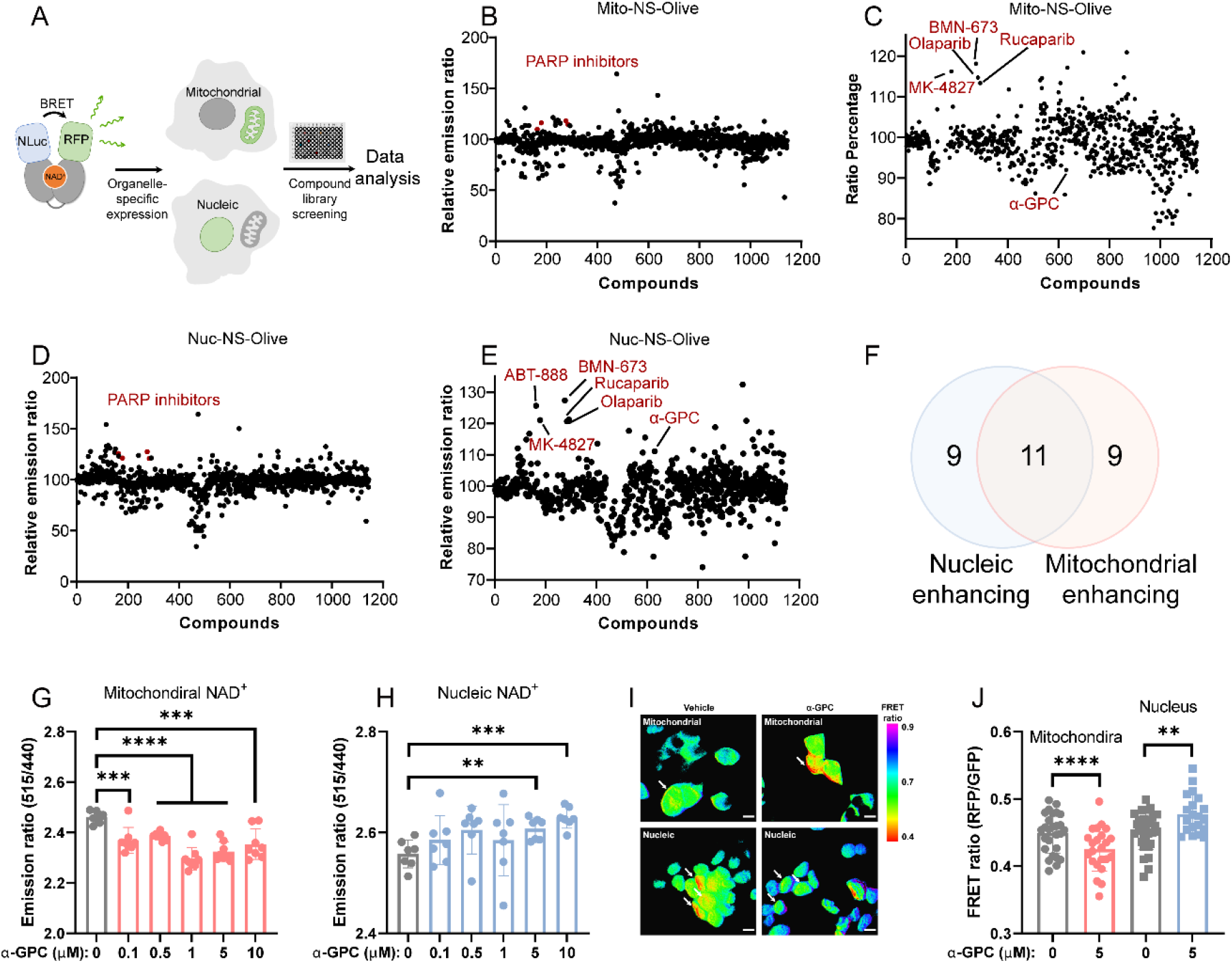
NS-Olive-based high-throughput screening for regulators of mitochondrial and nucleic NAD^+^. (A) Workflow of compound library screening. HEK 293T cells expressing organelle-specific NS-Olive were incubated with compounds for 24 h in white 96-well plates for BRET ratio measurement. (B) Effects of compounds on BRET ratios of cells expressing Mito-NS-Olive compared to non-treated control. (C) Remaining compounds that did not decrease Mito-NS-Olive luminescence to 70% of the control. (D) Effects of compounds on BRET ratios of cells expressing Nuc-NS-Olive compared to non-treated control. (E) Remaining compounds that did not decrease Nuc-NS-Olive luminescence to 70% of the control. (F) Venn diagram showing overlap of top 20 compounds that improve mitochondrial and nucleic NAD^+^. (G, H) Effects of α-GPC on subcellular NAD^+^ levels measured by Mito-NS-Olive and Nuc-NS-Olive, n ≥ 6. (I) Representative images of mitochondrial and nucleic FRET ratios of NS-Grapefruit in cells treated by 5 μM α-GPC. α-GPC decreased FRET in mitochondria but increased FRET in nucleus. (J) Summary of FRET ratio change of cells treated by 5 μM α-GPC. n ≥ 20. All statistical significance was calculated by Student’s t test. All data are represented as mean ± SD. *, *P* < 0.05; **, *P* < 0.01; ***, *P* < 0.001; ****, *P* < 0.0001.

Among the identified compounds, α-GPC notably decreases mitochondrial NAD^+^ but increases NAD^+^ in nucleus (Table S2 and S3). Additional tests with α-GPC at various concentrations confirmed the screening result (Fig. 4G-H). To visualize further this bidirectional effect on organelles under microscope, we established HEK 293T cell lines that express the FRET NAD^+^ sensor NS-Grapefruit in mitochondria (Mito-NS-Grapefruit) and nucleus (Nuc-NS-Grapefruit), which indicate NAD^+^ concentrations via FRET ratio measured at 515 nm and 595 nm. This reporter cell line distinguished the effects of FK866 and NRH (Fig. S6) as positive controls, demonstrating its proper function. The microscopic FRET image of the α-GPC-treated cells confirmed that α-GPC indeed increased the nucleic NAD^+^ but decreased NAD^+^ in mitochondria (Fig. 4I-J) relative to the total adenine nucleotide pool, making it a potential tool compound for redistributing subcellular NAD^+^.

## DISCUSSION

Quantifying and imaging cellular NAD^+^ dynamics requires more specific and pH-resistant NAD^+^ sensors. The previously reported sensors are susceptible to interferences induced by structurally similar analytes including NADH, NMN and NR^9,11^. In this work, we demonstrated that NS-Goji, Olive, and Grapefruit are specific towards NAD^+^ with little or no response towards these metabolites. Our sensors are based on Resonance Energy Transfer (RET) and avoided the use of circular permuted fluorescent proteins (cpFPs). The resulting sensors demonstrated a much-improved pH resistance^9,11,26^, which could be due to the use of structurally more stable fluorophores compared to cpFPs^27^. Another feature of the RET-based NAD^+^ sensors is the ratiometric readout that automatically corrects expression-related artifacts, making the quantification independent from calibrating fluorophores that are commonly needed for cpFP-based sensors^28,29^. Despite of the sensors’ improvement in NADH- and pH-resistance, our sensors are still interfered by high concentrations (millimolar range) of ATP and ADP. The physiological concentration of ATP is generally believed to be within the millimolar level^30,31^, and previous works also showed that the free intracellular ATP levels are in the micromolar range^32,33^. As only free, instead of bound, ATP and ADP potentially affect the sensors’ function, and the total adenine nucleotide pool remains constant in physiological conditions, the sensors’ response should hence reflect the dynamics of cellular NAD^+^ and not that of the potential interferents^9,12^. The modular RET sensor design allowed the signal output via both bioluminescence and fluorescence depending on the RET donors used, making the sensor design versatile for both high-throughput screening and microscopic imaging. Taking together, the proposed NAD^+^ sensors provided a set of multicolor, pH-stable, and analyte-specific tools for quantifying NAD^+^ in live cells and clinical samples.

Pharmacological modulation of NAD^+^ metabolism is a promising strategy for intervening various age-related diseases^34,35^. Moreover, the spatiotemporal distribution of NAD^+^ across the subcellular compartments integrate key signals in energy metabolism, gene transcription, and protein post-translational modifications in live cells^36,37^. However, the effects of different NAD^+^-modulating compounds and drugs on compartmentalized NAD^+^ are yet to be mapped. Here we showed that different NAD^+^ precursors and NAD^+^ metabolism-targeting drugs have distinct effects on cytosolic, nucleic, and mitochondrial NAD^+^. As dysregulated subcellular NAD^+^ dynamics play key roles in disease-related processes such as adipocyte differentiation^38^ and enhanced MARylation in ovarian cancer^39^, compounds’ effect on compartmentalized NAD^+^ should be considered during drug assessments and potentially leveraged as new strategies for drug design. Currently, NAD^+^-regulating strategies for improving age-related diseases primarily focus on increasing cellular NAD^+^. But considering the effects of shifted NAD^+^ compartmentalization during pathological events, the rebalancing of subcellular NAD^+^ could be an effective strategy for drug development. In our work, we provided a sensor-based procedure to screen for modulators of mitochondrial and nucleic NAD^+^ and identified that α-GPC shifts the subcellular NAD^+^ from mitochondria to nucleus. Future screenings of larger compound libraries should identify more potent and organelle-specific NAD^+^ modulators.

In summary, we developed BRET- and FRET-based sensors for quantifying NAD^+^ in samples ranging from lysed blood to organelles in live cells. The sensors will facilitate the assessment of NAD^+^ levels for clinical studies, as well as the elucidation of subcellular NAD^+^ dynamics for developing safer and more effective NAD^+^-modulating strategies.

## Methods

### Cloning

DNA sequences that encode mScarlet, *Ef*LigA, cpNluc and mNeoGreen were synthesized by Ruibiotech Co. (Guangzhou, China). Subdomains of these DNA sequences were amplified by PCR with 18 nt-overlapping ends to facilitate seamless cloning into pET51b vector for sensor expression in *E. coli* BL212 (DE3). His- and Strep-tags were used in the vector for the subsequent protein purification. For the sensor expression in mammalian cells, the entire sensor coding sequences were subcloned into pCDH-CMV-MCS-EF1-Neo vector. The mitochondrial targeting sequences (MTS) and nucleic targeting sequences (NTS) were added to the N termini of the sensor for mitochondrial and nucleic expression.

MTS:

ATGTCCGTCCTGACGCCGCTGCTGCTGCGGGGCTTGACAGGCTCGGCCCGGCGGCTCCCAGTGCCGCGCGCCAAGATCCATTCGTTG

NTS:

ATGGATCCAAAAAAGAAGAGAAAGGTA

### Mutagenesis

Sensor variants were generated by site-directed mutagenesis with specific or degenerate oligonucleotide primers purchased from Ruibiotech Co. Amplification was performed using KOD plus neo polymerase (Toyobo). More than 500 oligonucleotides were used; the sequences and resulting vector maps of the mutagenesis library are available upon request. Each variant was confirmed by sequencing.

### Protein expression and purification

All recombinant proteins were expressed in *E. coli* BL21 (DE3). The *E. coli* was cultured in 200 mL LB medium containing 50 μg/mL ampicillin at 37 °C until the cultures reached an OD^600^ of 0.6. Then, the growth temperature was shifted to 25 °C and protein expression was induced by adding 0.2 mM IPTG. After 16 h of incubation, bacteria were harvested and suspended in 50 mM HEPES buffer, pH 7.2, containing 500 mM NaCl and 10 mM imidazole, and lysed via high pressure homogenizer. Cell lysate was centrifuged at 15,000 × *g* for 15 min at 4 °C, and the supernatant was loaded sequentially onto Ni-NTA (Smart Lifesciences Inc., China) and Strep-Tactin columns (Smart Lifesciences Inc., China) for purification. The purified protein was desalted and exchanged into 50 mM HEPES buffer containing 50 mM NaCl (pH 7.2) with an Amicon Ultra-15 centrifugal filter (10 kDa MWCO, Merk Millipore Inc.). Protein was diluted to the required concentration before *in vitro* assays.

### Characterization of sensors *in vitro*

The sensors were titrated with NAD^+^, NADH, NADP^+^, NADPH, NMN and NRH to assess their sensitivity and selectivity. For BRET sensors, various concentrations of the compounds were mixed with 1 nM sensor and 1,000-fold diluted furimazine (Nano-Glo Luciferase Assay Substrate, Promega) in 100 μL of buffer (50 mM HEPES, 50 mM NaCl, pH 7.2) in a white 96-well plate (Grener). Bioluminescence was measured using a FlexStation® 3 Multi-Mode Microplate Reader (Molecular Devices) with NLuc emission measured at 440 nm and mScarlet-I emission at 590 nm, or mNeoGreen emission at 515 nm. For FRET sensors, various concentrations of the cofactors were mixed with 2 μM sensor in 100 μL of buffer (50 mM HEPES, 50 mM NaCl, pH 7.2) in a black clear bottom 96-well plates (Grener). Fluorescence was measured in a FlexStation® 3 Multi-Mode Microplate Reader (Molecular Devices) with excitation measured at 470 nm and emission at 515 nm and 590 nm. The ratios between NLuc and mScarlet-I emission (mScarlet-I/NLuc), and mNeoGreen (mNeoGreen/NLuc), or ratios between mScarlet-I and mNeoGreen (mScarlet-I/mNeoGreen) were plotted against the compound concentrations.

### Generation and screening of protein library

To generate sensor mutant libraries, overlapping PCR and oligonucleotides containing degenerate codons (VST) were used to randomize amino acid residues on the linker regions of the sensor. Then, *E*.*coli* BL21(DE3) was transformed with the plasmid library and spread out on LB agar plates containing 100 μg/mL ampicillin. After overnight incubation, single colonies were inoculated into 96-well plates containing 200 μL LB medium with 100 μg/mL ampicillin. The bacterial cultures were incubated overnight at 37 °C. Then, 100 μL of the culture were transferred to 2 mL 96-deep-well plates containing 900 μL LB medium with 100 μg/mL ampicillin and 0.2 mM IPTG. The rest of the culture was stored at 4 °C. After a 16 h-incubation at 25 °C with 280 rpm shaking rate, the bacteria were collected by centrifugation at 4,000 *g* for 10 min. Cell lysates were then prepared from bacterial pellets using a buffer solution containing 50 mM HEPES, 50 mM NaCl, 0.1 mg/mL chicken egg lysozyme and 0.2 U/mL Benzonase (pH 7.2). Crude lysate was centrifuged at 4,000 *g* for 30 min. 100 μL of supernatant was transferred into flat-bottom black 96-well plates (Greiner). Initial fluorescence was measured in a FlexStation® 3 Multi-Mode Microplate Reader (Molecular Devices) with excitation at 470 nm and emission at 515 nm and 590 nm. Fluorescence was assayed again after the addition of 10 μL 10 mM NAD^+^. After calculating the Ratio (590/515) before and after the addition of NAD^+^, plasmids were amplified for promising candidates and were subject to subsequent rounds of mutagenesis.

### In vitro measurement of NAD^+^ using NS-Goji 1.3 and LC-MS

Blood samples were lysed by adding 4 volumes of 0.5 N perchloric acid into 1 volume of sample. The resulting mixture was thoroughly vortexed for 2 min. Then, the lysate was centrifuged at 12,000 *g*, 4 °C for 5 min. The supernatant was then diluted 10-fold in 10 × HEPES buffer (500 mM NaCl, 500 mM HEPES, pH 7.2). Finally, the resulting supernatant was diluted 10-fold in the assay buffer (50 mM NaCl, 50 mM HEPES, pH 7.2) containing 1 nM sensor NS-Goji 1.3 and 1,000-fold diluted furimazine. Bioluminescence was measured using a FlexStation® 3 Multi-Mode Microplate Reader (Molecular Devices) with NLuc emission measured at 440 nm and mScarlet-I at 590 nm. The titration was performed by measuring the emission ratios (R) of known concentrations of NAD^+^. The sensor’s maximum ratio (R_max_), minimum ratio (R_min_), c50 and Hill coefficient (h) were obtained by fitting the data to the Hill-Langmuir equation.

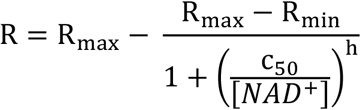

NAD^+^ concentrations in unknown samples were calculated with the rearranged the equation using the measured R.

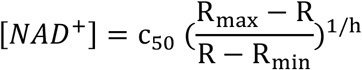

In parallel, the samples were analyzed by HPLC-MS use Agilent InfinityLab LC/MSD iQ 6160A. Freshly prepared sample (6 μL) was injected on an ZORBAX SB-Aq column (Agilent, 4.6 × 100 mm, 3.5 μm). The column temperature was set to 45 °C. The analytes were eluted using a 1 mL/min flow of an aqueous acetonitrile (A) and 0.1% formic acid-methyl alcohol (B). After sample injection, a 3 min organic phase gradient was applied as following: 0 min 20% A, 1.2 min 60% A, 2.5 min 60% A, 2.6 min 20% A, 3 min 20% A. Subsequently, the column was washed using 100% acetonitrile and re-equilibrated. NAD^+^ featured a retention time of 1.1 min. Negative polarity electrospray ionization was selected for MS analysis. Samples were ionized under high-flow conditions with curtain gas = 35 psi, collision gas = “high”, capillary voltage = 3500 V, and dry temperature = 325 °C. The following ion pairs were used for analyte identification: NAD^+^1, 662 → 539.7 m/z; NAD^+^2, 662 → 80 m/z. NAD^+^ concentrations in the standard curve were 0.0500, 0.100, 0.200, 0.500, and 1.00 μg/mL. A linear regression was then applied (weighting 1/x, r=0.9994) to obtain the standard curve. With this method, the limit of detection (LOD) was 0.0200 μg/mL (Signal/Noise ≥ 3) and limit of quantification (LOQ) was 0.0500 μg/mL (Signal/Noise ≥ 10).

### Cell culture, lentivirus preparation and infection

HEK 293T cells were cultured in DMEM (11885, Gibco) supplemented with 10% FBS (Gibco 10082147) at 37 °C with 5% CO_2_. Plasmids pMD2G, psPAX2 and pCDH-CMV-NADS-EF1-Neo were used for lentivirus production. The viral supernatant was filtered through a 0.45 μm PVDF filter and further concentrated by PEG-8000 precipitation. The resulting viral solution was aliquoted, flash frozen and stored at −80 °C. HEK 293T cells cultured in 60 mm plates were infected with virus solutions and the culture media was changed 6 h after infection. 48 h after the infection, the cells were harvested and serial-diluted to select infected cells with sensor’s fluorescence as selection marker.

### Cellular BRET and FRET measurement using microplate reader

HEK 293T cell line that stably expresses the BRET sensor was seeded at a density of 10,000 cells per well in a white 96-well plate (Grener) with DMEM culture (11885, Gibco) supplemented with 10% FBS (Gibco 10082147). After 24 h of incubation at 37 °C with 5% CO_2_, different compounds were added into the culture medium. After 24 h of treatment, culture medium was removed by aspiration and replaced with 100 μL phenol red-free DMEM culture containing 1,000-fold diluted furimazine. Bioluminescence was measured using a FlexStation® 3 Multi-Mode Microplate Reader (Molecular Devices) with NLuc emission measured at 440 nm and mScarlet-I emission at 590 nm.

For FRET measurement in live cells, FRET sensor-expressing HEK 293T cell line was seed at a density of 10,000 cells per well in a black clear bottom plate (Grener) with DMEM culture (11885, Gibco) supplemented with 10% FBS (Gibco 10082147). After 48 h incubation at 37 °C with 5% CO_2_, the fluorescence was measured in a FlexStation® 3 Multi-Mode Microplate Reader (Molecular Devices) with excitation at 470 nm and emission at 515 nm and 590 nm.

### Imaging

The imaging was performed with an Olympus IX83 inverted microscope equipped with 100 × oil immersion objective (N.A. 1.50). FRET imaging was performed with excitation at 488 nm, emissions at 532 nm and 561 nm. All analyses were performed using ImageJ v1.53K. The region of interest (ROI) was selected from single cells. The background was subtracted from each emission channel and the resulting 561 nm / 532 nm ratio was used to indicate NAD^+^ levels.

**Correspondence and requests for materials** should be address to M.H. and Q.Y.

## ACKNOWLEDGEMENT

This work is supported by Shenzhen Science and Technology Program (ZDSYS20210623091810032), National Key R&D Program of China (2020YFC2002904, 2021YFF1200302), and Shenzhen Institute of Advanced Technology, Chinese Academy of Sciences.

## AUTHOR CONTRIBUTIONS

Q.Y. and L.C. conceived this study. L.C. and Q.Y. designed the sensors. M.C. performed HPLC-MS quantifications. Y.L. and L.C. performed cell experiments. M.L performed chemical synthesis. M.H. and B.L. provided clinical samples. All authors contributed to data analysis or manuscript writing.

## Reference

(1) Covarrubias, A. J.; Perrone, R.; Grozio, A.; Verdin, E. NAD+ Metabolism and Its Roles in Cellular Processes during Ageing. Nat Rev Mol Cell Biol 2021, 22 (2), 119–141. https://doi.org/10.1038/s41580-020-00313-x.

(2) Shi, H.; Enriquez, A.; Rapadas, M.; Martin, E. M. M. A.; Wang, R.; Moreau, J.; Lim, C. K.; Szot, J. O.; Ip, E.; Hughes, J. N.; Sugimoto, K.; Humphreys, D. T.; McInerney-Leo, A. M.; Leo, P. J.; Maghzal, G. J.; Halliday, J.; Smith, J.; Colley, A.; Mark, P. R.; Collins, F.; Sillence, D. O.; Winlaw, D. S.; Ho, J. W. K.; Guillemin, G. J.; Brown, M. A.; Kikuchi, K.; Thomas, P. Q.; Stocker, R.; Giannoulatou, E.; Chapman, G.; Duncan, E. L.; Sparrow, D. B.; Dunwoodie, S. L. NAD Deficiency, Congenital Malformations, and Niacin Supplementation. New England Journal of Medicine 2017, 377 (6), 544–552. https://doi.org/10.1056/NEJMoa1616361.

(3) Lautrup, S.; Sinclair, D. A.; Mattson, M. P.; Fang, E. F. NAD+ in Brain Aging and Neurodegenerative Disorders. Cell Metabolism 2019, 30 (4), 630–655. https://doi.org/10.1016/j.cmet.2019.09.001.

(4) Chini, E. N. Of Mice and Men: NAD+ Boosting with Niacin Provides Hope for Mitochondrial Myopathy Patients. Cell Metabolism 2020, 31 (6), 1041–1043. https://doi.org/10.1016/j.cmet.2020.05.013.

(5) Sciarretta, S.; Forte, M.; Castoldi, F.; Frati, G.; Versaci, F.; Sadoshima, J.; Kroemer, G.; Maiuri, M. C. Caloric Restriction Mimetics for the Treatment of Cardiovascular Diseases. Cardiovasc Res 2021, 117 (6), 1434–1449. https://doi.org/10.1093/cvr/cvaa297.

(6) Yoshino, J.; Baur, J. A.; Imai, S. Nad+ Intermediates: The Biology and Therapeutic Potential of NMN and NR. Cell Metabolism 2018, 27 (3), 513–528. https://doi.org/10.1016/j.cmet.2017.11.002.

(7) Janssens, G. E.; Grevendonk, L.; Perez, R. Z.; Schomakers, B. V.; de Vogel-van den Bosch, J.; Geurts, J. M. W.; van Weeghel, M.; Schrauwen, P.; Houtkooper, R. H.; Hoeks, J. Healthy Aging and Muscle Function Are Positively Associated with NAD+ Abundance in Humans. Nat Aging 2022, 1–10. https://doi.org/10.1038/s43587-022-00174-3.

(8) Yoshino, M.; Yoshino, J.; Kayser, B. D.; Patti, G. J.; Franczyk, M. P.; Mills, K. F.; Sindelar, M.; Pietka, T.; Patterson, B. W.; Imai, S.-I.; Klein, S. Nicotinamide Mononucleotide Increases Muscle Insulin Sensitivity in Prediabetic Women. Science 2021, 372 (6547), 1224–1229. https://doi.org/10.1126/science.abe9985.

(9) Zou, Y.; Wang, A.; Huang, L.; Zhu, X.; Hu, Q.; Zhang, Y.; Chen, X.; Li, F.; Wang, Q.; Wang, H.; Liu, R.; Zuo, F.; Li, T.; Yao, J.; Qian, Y.; Shi, M.; Yue, X.; Chen, W.; Zhang, Z.; Wang, C.; Zhou, Y.; Zhu, L.; Ju, Z.; Loscalzo, J.; Yang, Y.; Zhao, Y. Illuminating NAD+ Metabolism in Live Cells and In Vivo Using a Genetically Encoded Fluorescent Sensor. Developmental Cell 2020, 53 (2), 240-252.e7. https://doi.org/10.1016/j.devcel.2020.02.017.

(10) Yu, Q.; Pourmandi, N.; Xue, L.; Gondrand, C.; Fabritz, S.; Bardy, D.; Patiny, L.; Katsyuba, E.; Auwerx, J.; Johnsson, K. A Biosensor for Measuring NAD+ Levels at the Point of Care. Nat Metab 2019, 1 (12), 1219–1225. https://doi.org/10.1038/s42255-019-0151-7.

(11) Cambronne, X. A.; Stewart, M. L.; Kim, D.; Jones-Brunette, A. M.; Morgan, R. K.; Farrens, D. L.; Cohen, M. S.; Goodman, R. H. Biosensor Reveals Multiple Sources for Mitochondrial NAD+. 2016, 5. https://doi.org/10/ghbc89.

(12) Chen, W.; Liu, S.; Yang, Y.; Zhang, Z.; Zhao, Y. Spatiotemporal Monitoring of NAD+ Metabolism with Fluorescent Biosensors. Mechanisms of Ageing and Development 2022, 111657. https://doi.org/10.1016/j.mad.2022.111657.

(13) Zhao, Y.; Jin, J.; Hu, Q.; Zhou, H.-M.; Yi, J.; Yu, Z.; Xu, L.; Wang, X.; Yang, Y.; Loscalzo, J. Genetically Encoded Fluorescent Sensors for Intracellular NADH Detection. Cell Metabolism 2011, 14 (4), 555–566. https://doi.org/10.1016/j.cmet.2011.09.004.

(14) Gajiwala, K. S.; Pinko, C. Structural Rearrangement Accompanying NAD+ Synthesis within a Bacterial DNA Ligase Crystal. Structure 2004, 12 (8), 1449–1459. https://doi.org/10.1016/j.str.2004.05.017.

(15) Gilmour, B. C.; Gudmundsrud, R.; Frank, J.; Hov, A.; Lautrup, S.; Aman, Y.; Røsjø, H.; Brenner, C.; Ziegler, M.; Tysnes, O.-B.; Tzoulis, C.; Omland, T.; Søraas, A.; Holmøy, T.; Bergersen, L. H.; Storm-Mathisen, J.; Nilsen, H.; Fang, E. F. Targeting NAD+ in Translational Research to Relieve Diseases and Conditions of Metabolic Stress and Ageing. Mechanisms of Ageing and Development 2020, 186, 111208. https://doi.org/10.1016/j.mad.2020.111208.

(16) Pérez, M. J.; Baden, P.; Deleidi, M. Progresses in Both Basic Research and Clinical Trials of NAD+ in Parkinson’s Disease. Mechanisms of Ageing and Development 2021, 197, 111499. https://doi.org/10.1016/j.mad.2021.111499.

(17) Klimova, N.; Kristian, T. Multi-Targeted Effect of Nicotinamide Mononucleotide on Brain Bioenergetic Metabolism. Neurochem Res 2019, 44 (10), 2280–2287. https://doi.org/10.1007/s11064-019-02729-0.

(18) Wang, G.; Han, T.; Nijhawan, D.; Theodoropoulos, P.; Naidoo, J.; Yadavalli, S.; Mirzaei, H.; Pieper, A. A.; Ready, J. M.; McKnight, S. L. P7C3 Neuroprotective Chemicals Function by Activating the Rate-Limiting Enzyme in NAD Salvage. Cell 2014, 158 (6), 1324–1334. https://doi.org/10.1016/j.cell.2014.07.040.

(19) Yao, H.; Liu, M.; Wang, L.; Zu, Y.; Wu, C.; Li, C.; Zhang, R.; Lu, H.; Li, F.; Xi, S.; Chen, S.; Gu, X.; Liu, T.; Cai, J.; Wang, S.; Yang, M.; Xing, G.-G.; Xiong, W.; Hua, L.; Tang, Y.; Wang, G. Discovery of Small-Molecule Activators of Nicotinamide Phosphoribosyltransferase (NAMPT) and Their Preclinical Neuroprotective Activity. Cell Res 2022, 1–15. https://doi.org/10.1038/s41422-022-00651-9.

(20) Nikiforov, A.; Kulikova, V.; Ziegler, M. The Human NAD Metabolome: Functions, Metabolism and Compartmentalization. Critical Reviews in Biochemistry and Molecular Biology 2015, 50 (4), 284–297. https://doi.org/10.3109/10409238.2015.1028612.

(21) Napper, A. D.; Hixon, J.; McDonagh, T.; Keavey, K.; Pons, J.-F.; Barker, J.; Yau, W. T.; Amouzegh, P.; Flegg, A.; Hamelin, E.; Thomas, R. J.; Kates, M.; Jones, S.; Navia, M. A.; Saunders, J. O.; DiStefano, P. S.; Curtis, R. Discovery of Indoles as Potent and Selective Inhibitors of the Deacetylase SIRT1. J. Med. Chem. 2005, 48 (25), 8045–8054. https://doi.org/10.1021/jm050522v.

(22) Shoba, B.; Lwin, Z. M.; Ling, L. S.; Bay, B.-H.; Yip, G. W.; Kumar, S. D. Function of Sirtuins in Biological Tissues. The Anatomical Record 2009, 292 (4), 536–543. https://doi.org/10.1002/ar.20875.

(23) Oplustil O’Connor, L.; Rulten, S. L.; Cranston, A. N.; Odedra, R.; Brown, H.; Jaspers, J. E.; Jones, L.; Knights, C.; Evers, B.; Ting, A.; Bradbury, R. H.; Pajic, M.; Rottenberg, S.; Jonkers, J.; Rudge, D.; Martin, N. M. B.; Caldecott, K. W.; Lau, A.; O’Connor, M. J. The PARP Inhibitor AZD2461 Provides Insights into the Role of PARP3 Inhibition for Both Synthetic Lethality and Tolerability with Chemotherapy in Preclinical Models. Cancer Res 2016, 76 (20), 6084–6094. https://doi.org/10.1158/0008-5472.CAN-15-3240.

(24) Javed, Z.; Sadia, H.; Iqbal, M. J.; Shamas, S.; Malik, K.; Ahmed, R.; Raza, S.; Butnariu, M.; Cruz-Martins, N.; Sharifi-Rad, J. Apigenin Role as Cell-Signaling Pathways Modulator: Implications in Cancer Prevention and Treatment. Cancer Cell Int 2021, 21 (1), 189. https://doi.org/10.1186/s12935-021-01888-x.

(25) Gardell, S. J.; Hopf, M.; Khan, A.; Dispagna, M.; Hampton Sessions, E.; Falter, R.; Kapoor, N.; Brooks, J.; Culver, J.; Petucci, C.; Ma, C.-T.; Cohen, S. E.; Tanaka, J.; Burgos, E. S.; Hirschi, J. S.; Smith, S. R.; Sergienko, E.; Pinkerton, A. B. Boosting NAD+ with a Small Molecule That Activates NAMPT. Nat Commun 2019, 10 (1), 3241. https://doi.org/10.1038/s41467-019-11078-z.

(26) Berg, J.; Hung, Y. P.; Yellen, G. A Genetically Encoded Fluorescent Reporter of ATP:ADP Ratio. Nat Methods 2009, 6 (2), 161–166. https://doi.org/10.1038/nmeth.1288.

(27) Kostyuk, A. I.; Demidovich, A. D.; Kotova, D. A.; Belousov, V. V.; Bilan, D. S. Circularly Permuted Fluorescent Protein-Based Indicators: History, Principles, and Classification. Int J Mol Sci 2019, 20 (17), 4200. https://doi.org/10.3390/ijms20174200.

(28) Su, S.; Phua, S. C.; DeRose, R.; Chiba, S.; Narita, K.; Kalugin, P. N.; Katada, T.; Kontani, K.; Takeda, S.; Inoue, T. Genetically Encoded Calcium Indicator Illuminates Calcium Dynamics in Primary Cilia. Nat Methods 2013, 10 (11), 1105–1107. https://doi.org/10.1038/nmeth.2647.

(29) Ast, C.; Foret, J.; Oltrogge, L. M.; De Michele, R.; Kleist, T. J.; Ho, C.-H.; Frommer, W. B. Ratiometric Matryoshka Biosensors from a Nested Cassette of Green- and Orange-Emitting Fluorescent Proteins. Nat Commun 2017, 8 (1), 431. https://doi.org/10.1038/s41467-017-00400-2.

(30) Greiner, J. V.; Glonek, T. Intracellular ATP Concentration and Implication for Cellular Evolution. Biology 2021, 10 (11), 1166. https://doi.org/10.3390/biology10111166.

(31) Patel, A.; Malinovska, L.; Saha, S.; Wang, J.; Alberti, S.; Krishnan, Y.; Hyman, A. A. ATP as a Biological Hydrotrope. Science 2017, 356 (6339), 753–756. https://doi.org/10.1126/science.aaf6846.

(32) Gajewski, C. D.; Yang, L.; Schon, E. A.; Manfredi, G. New Insights into the Bioenergetics of Mitochondrial Disorders Using Intracellular ATP Reporters. MBoC 2003, 14 (9), 3628–3635. https://doi.org/10.1091/mbc.e02-12-0796.

(33) Kim, J.-H.; Ahn, J.-H.; Barone, P. W.; Jin, H.; Zhang, J.; Heller, D. A.; Strano, M. S. A Luciferase/Single-Walled Carbon Nanotube Conjugate for Near-Infrared Fluorescent Detection of Cellular ATP. Angewandte Chemie 2010, 122 (8), 1498–1501. https://doi.org/10.1002/ange.200906251.

(34) Yoshino, J.; Mills, K. F.; Yoon, M. J.; Imai, S. Nicotinamide Mononucleotide, a Key NAD+ Intermediate, Treats the Pathophysiology of Diet- and Age-Induced Diabetes in Mice. Cell Metabolism 2011, 14 (4), 528–536. https://doi.org/10.1016/j.cmet.2011.08.014.

(35) Mills, K. F.; Yoshida, S.; Stein, L. R.; Grozio, A.; Kubota, S.; Sasaki, Y.; Redpath, P.; Migaud, M. E.; Apte, R. S.; Uchida, K.; Yoshino, J.; Imai, S. Long-Term Administration of Nicotinamide Mononucleotide Mitigates Age-Associated Physiological Decline in Mice. Cell Metabolism 2016, 24 (6), 795–806. https://doi.org/10.1016/j.cmet.2016.09.013.

(36) Cambronne, X. A.; Kraus, W. L. Location, Location, Location: Compartmentalization of NAD+ Synthesis and Functions in Mammalian Cells. Trends in Biochemical Sciences 2020, 45 (10), 858–873. https://doi.org/10.1016/j.tibs.2020.05.010.

(37) Zhu, Y.; Liu, J.; Park, J.; Rai, P.; Zhai, R. G. Subcellular Compartmentalization of NAD+ and Its Role in Cancer: A SereNADe of Metabolic Melodies. Pharmacol Ther 2019, 200, 27–41. https://doi.org/10.1016/j.pharmthera.2019.04.002.

(38) Ryu, K. W.; Nandu, T.; Kim, J.; Challa, S.; DeBerardinis, R. J.; Kraus, W. L. Metabolic Regulation of Transcription through Compartmentalized NAD+ Biosynthesis. Science 2018, 360 (6389), eaan5780. https://doi.org/10.1126/science.aan5780.

(39) Challa, S.; Khulpateea, B. R.; Nandu, T.; Camacho, C. V.; Ryu, K. W.; Chen, H.; Peng, Y.; Lea, J. S.; Kraus, W. L. Ribosome ADP-Ribosylation Inhibits Translation and Maintains Proteostasis in Cancers. Cell 2021, 184 (17), 4531-4546.e26. https://doi.org/10.1016/j.cell.2021.07.005.

