## supplemental file for "Ratiometric NAD^+^ sensors reveal subcellular NAD^+^ modulators"

**Affiliations:**

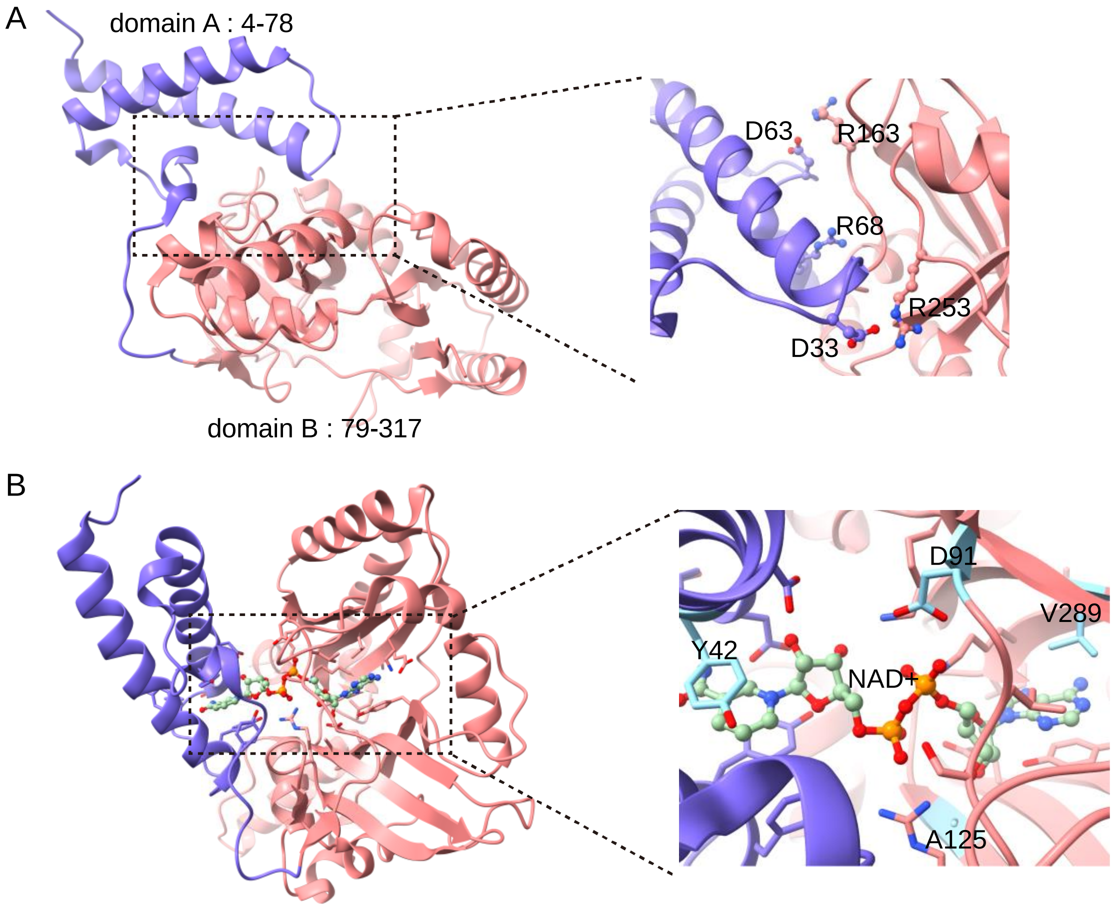


**Fig.S1 Structure analysis of *Ef*LigA.** (A) Structure of *Ef*LigA (PDB ID: 1ta8) with domain A (residues 4-78) colored in blue and domain B (residues 79-317) in pink. Residues potentially contributing to the domain interactions (D33, D63, R68, R163, and R253) are shown in ball stick. (B) Structure of *Ef*LigA bound to native ligand $NAD^+$ (PDB ID: 1tae). $NAD^+$ is shown in ball stick and potential interacting residues (Y42, D91, A125, and V289) shown in stick.

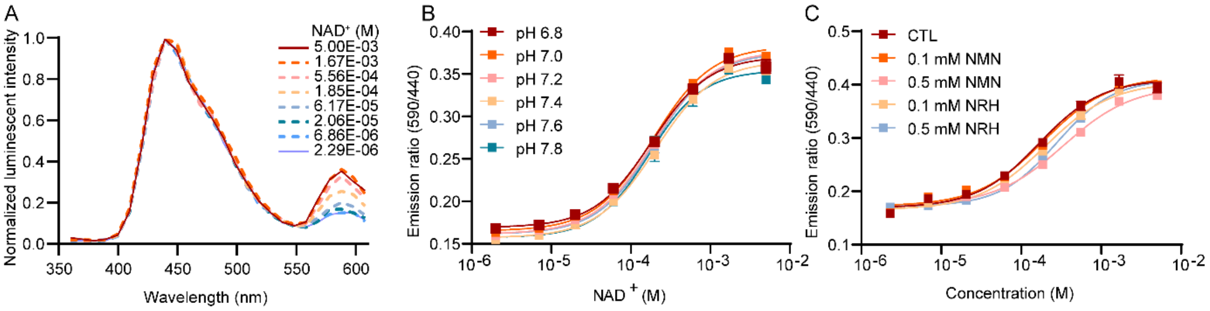


**Fig.S2 Performance of NS-Goji 1.0.** (A) Emission spectra of NS-Goji 1.0 at various $NAD^+$ concentrations. (B) Titration curve of NS-Goji 1.0 at pH 6.8-7.8. The emission ratio of NS-Goji 1.0 is minimally affected by changes in pH. (C) Titration curve of NS-Goji 1.0 against $NAD^+$ in the presence of 0.1 mM or 0.5 mM NMN and 0.1 mM or 0.5 mM NRH.

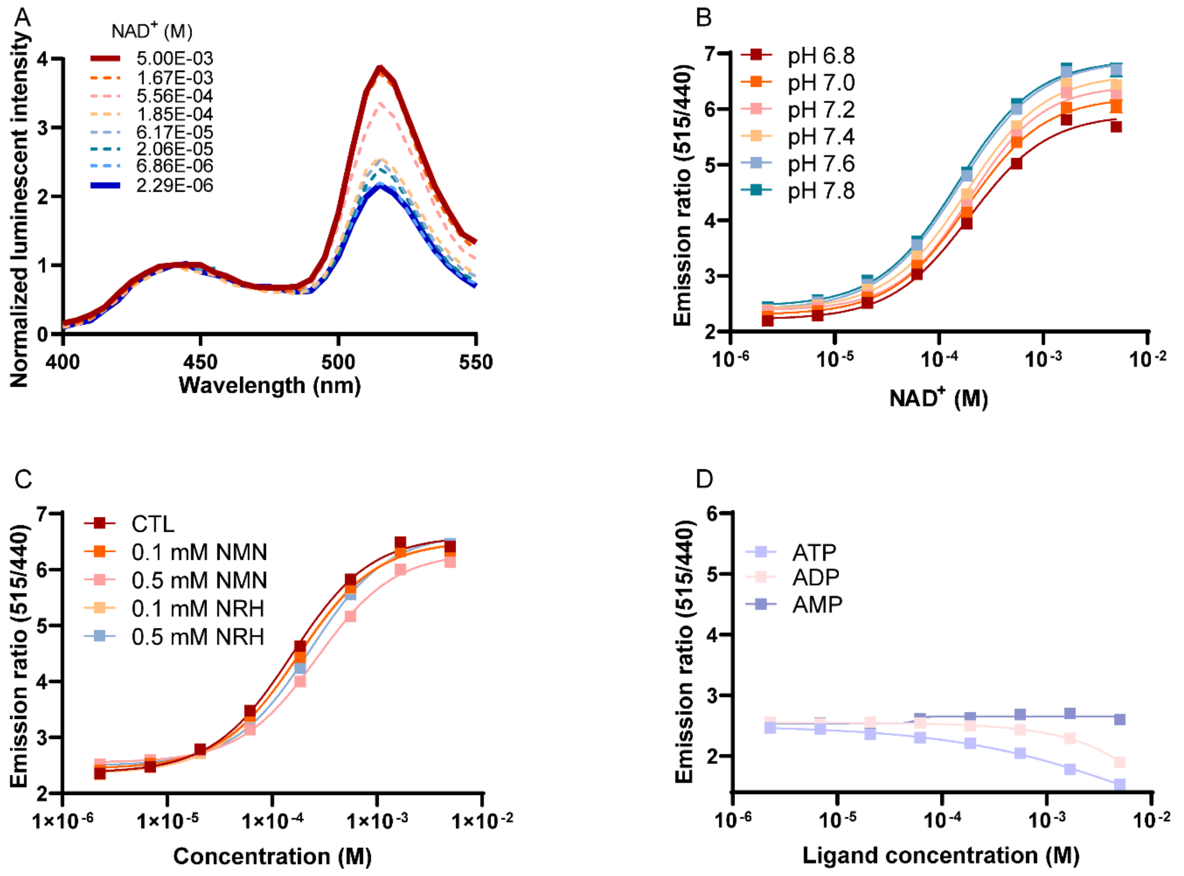


**Fig.S3 Performance of NS-Olive**. (A) Emission spectra of NS-Olive at various $NAD^+$ concentrations. (B) Titration curve of NS-Olive at pH 6.8-7.8. The emission ratio of NS-Olive is slightly affected by changes in pH. (C) Titration curve of NS-Olive against $NAD^+$ in the presence of 0.1 mM or 0.5 mM NMN and 0.1 mM or 0.5 mM NRH. (D) Titration curve of NS-Olive with ATP, ADP, and AMP.

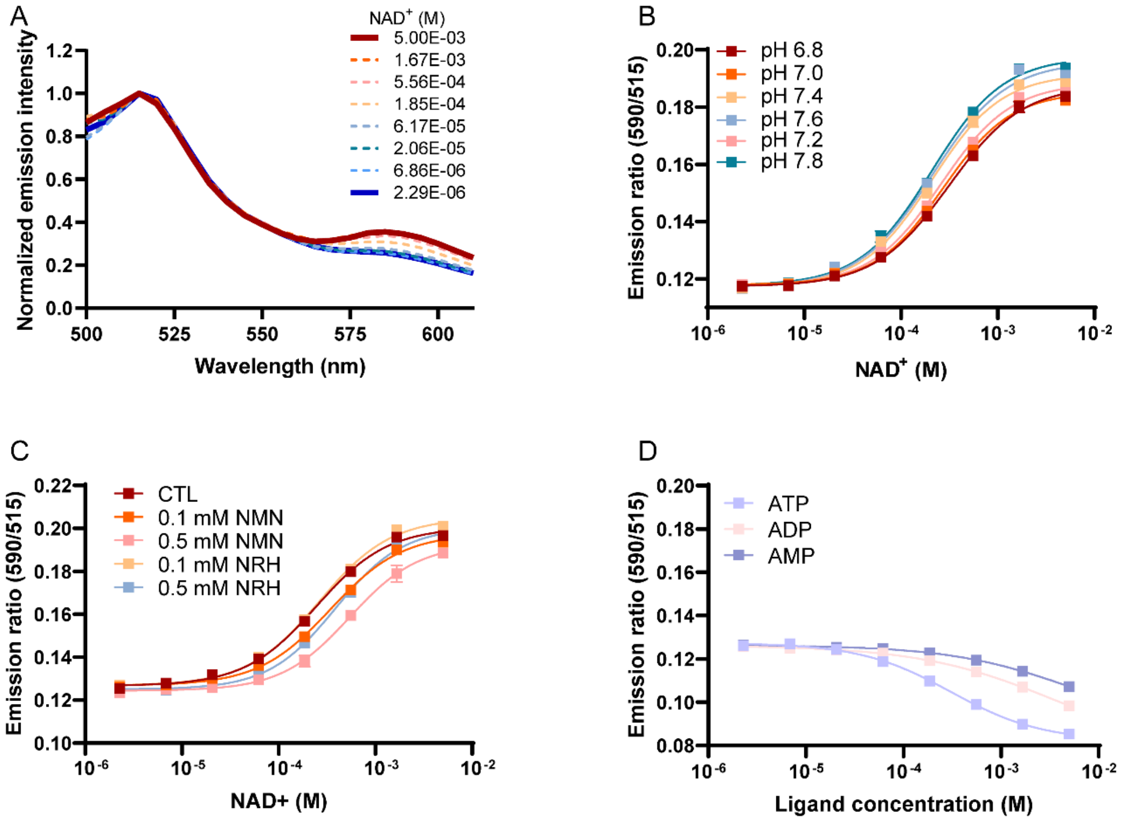


**Fig.S4 Performance of NS-Grapefruit**. (A) Emission spectra of NS-Grapefruit at various $NAD^+$ concentrations. (B) Titration curve of NS-Grapefruit at pH 6.8-7.8. The emission ratio of NS-Grapefruit is slightly affected by changes in pH. (C) Titration curve of NS-Grapefruit against $NAD^+$ in the presence of 0.1 mM or 0.5 mM NMN and 0.1 mM or 0.5 mM NRH. (D) Titration curve of NS-Grapefruit with ATP, ADP, and AMP.

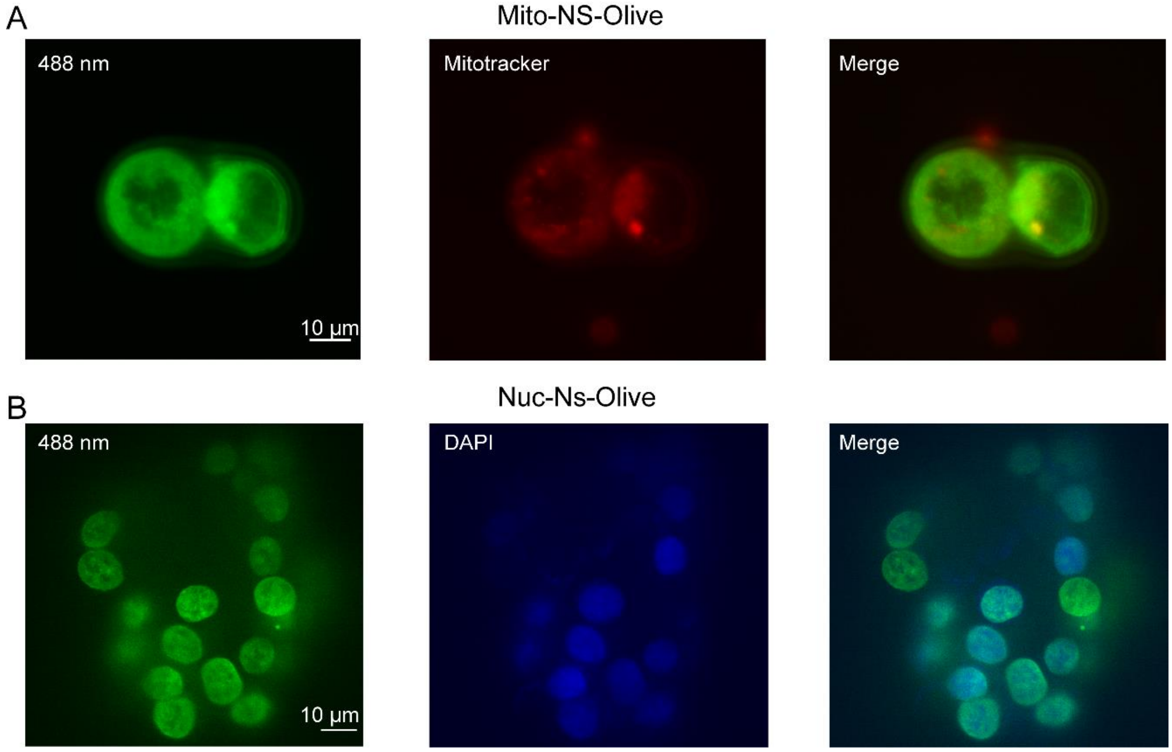


**Fig.S5 Representative microscopic images of cells stably express NS-Olive** in (A) mitochondria and (B) nucleus.

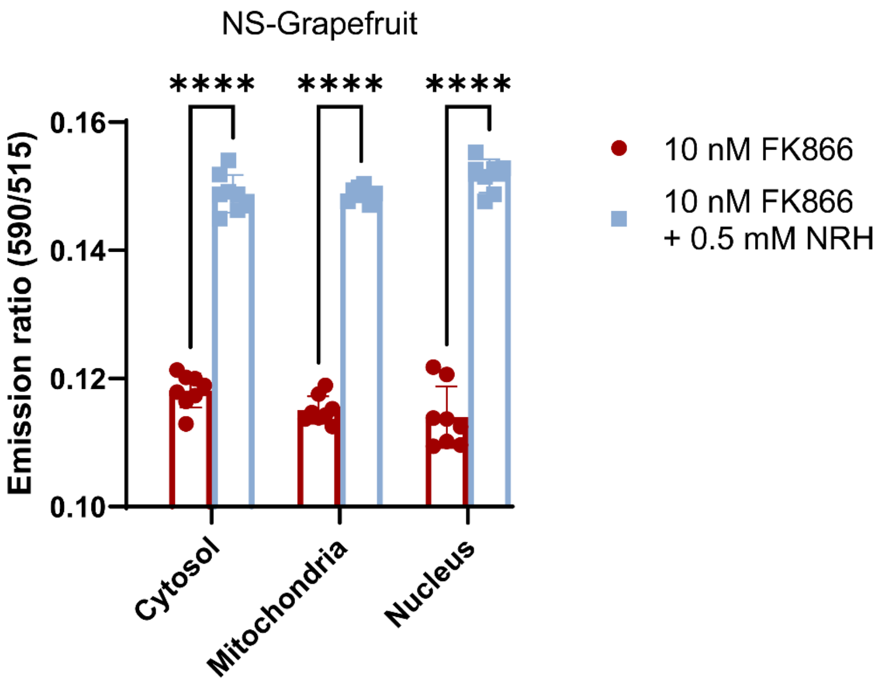


**Fig.S6 Cellular performance of NS-Grapefruit measured by microplate reader.**

Emission ratio of 590 nm and 515 nm were used as indicator of cellular $NAD^+$. HEK 293T cells were treated by 10 nM FK866 with and without 0.5 mM NRH for 24 h.

**Table S1. Summary of $NAD^+$ sensor performance.**

| Sensors | $R_{max}$ | $R_{min}$ | $R_{max}/R_{min}$ | c50 |
|---|---|---|---|---|
| NS-Goji 1.0 | 0.38 | 0.17 | 2.2 | 235.7 ± 7.9 μM |
| NS-Goji 1.1 | 0.30 | 0.16 | 1.8 | 3.1 ± 0.3 mM |
| NS-Goji 1.2 | 0.64 | 0.19 | 3.3 | 7.0 ± 0.1 μM |
| NS-Goji 1.3 | 0.50 | 0.16 | 3.1 | 47.4 ± 0.9 nM |
| NS-Olive | 5.99 | 2.25 | 2.6 | 254.7 ± 11.5 μM |
| NS-Grapefruit | 0.20 | 0.12 | 1.6 | 244.2 ± 14.9 μM |

**Table S2. Top 20 compounds for increasing $NAD^+$ in nucleus.**

| No. | Compounds | Relative ratio of Nuc-NS-Olive (%) | Relative luminescence | Targets |
|---|---|---|---|---|
| 1 | BMN 673* | 127.4 | 101% | PARP1 |
| 2 | ABT-888 | 125.7 | 92% | PARP1/2 |
| 3 | Rucaparib* | 121.3 | 130% | PARP1 |
| 4 | MK-4827* | 121.1 | 93% | PARP1/2 |
| 5 | Phenazopyridine HCl | 120.9 | 142% | Sodium channel |
| 6 | Olaparib* | 120.7 | 104% | PARP1/2 |
| 7 | Raloxifene HCl* | 118.9 | 105% | Estrogen receptor |
| 8 | Dinaciclib* | 117.7 | 141% | CDK2/5/1/9 |
| 9 | Triclabendazole | 117.1 | 106% | Tubulin |
| 10 | Ruxolitinib | 114.9 | 183% | JAK2/1 |
| 11 | TSU-68* | 113.6 | 106% | PDGFRβ |
| 12 | Econazole nitrate | 112.9 | 137% | Calcium channel |
| 13 | Pimecrolimus | 112.7 | 143% | - |
| 14 | Azaguanine-8* | 112.5 | 107% | - |
| 15 | Doxazosin mesylate | 112.1 | 94% | α1-adrenergic receptor |
| 16 | Bifonazole* | 111.9 | 186% | Aromatase |
| 17 | Alpha-GPC | 111.1 | 189% | - |
| 18 | Clomiphene citrate* | 110.8 | 90% | Estrogen receptor |
| 19 | Thiamphenicol | 110.6 | 85% | Anti-infection |
| 20 | Triflurdine* | 108.2 | 72% | DNA synthesis |

"*" compound appeared in both screenings

"-" unclear.

**Table S3. Top 20 compounds for increasing $NAD^+$ in nucleus.**

| No. | Compounds | Relative ratio of Mito-NS-Olive (%) | Relative luminescence | Targets |
|---|---|---|---|---|
| 1 | Triflurdine* | 120.9 | 91% | DNA synthesis |
| 2 | Raloxifene HCl* | 120.9 | 108% | Estrogen receptor |
| 3 | BMN 673* | 118.1 | 117% | PARP1 |
| 4 | Oxaliplatin | 117.1 | 131% | DNA synthesis |
| 5 | MK-4827* | 116.2 | 99% | PARP1/2 |
| 6 | Procarbazine HCl | 114.7 | 117% | DNA synthesis |
| 7 | Olaparib* | 114.7 | 107% | PARP1/2 |
| 8 | Pasiniazid | 114.5 | 129% | - |
| 9 | Dinaciclib* | 114.1 | 119% | CDK2/5/1/9 |
| 10 | Rucaparib* | 113.3 | 152% | PARP1 |
| 11 | Rifaximin | 113.3 | 126% | RNA polymerase |
| 12 | KPT-330 | 113.0 | 78% | CRM1 |
| 13 | Nicotine ditartrate | 112.1 | 126% | - |
| 14 | TSU-68* | 111.8 | 78% | PDGFRβ |
| 15 | Sulfadiazine | 111.1 | 126% | Antibiotic |
| 16 | Mizoribine | 110.6 | 107% | Monophosphate synthetase |
| 17 | Clomiphene citrate* | 109.5 | 80% | Estrogen receptor |
| 18 | Azaguanine-8* | 109.4 | 93% | - |
| 19 | Linifanib | 106.9 | 75% | VEGFR/PDGFR |
| 20 | Bifonazole* | 102.5 | 105% | Aromatase |

"*" compound appeared in both screenings

"-" unclear.
